# Neural correlates of the DMT experience as assessed via multivariate EEG

**DOI:** 10.1101/706283

**Authors:** Christopher Timmermann, Leor Roseman, Michael Schartner, Raphael Milliere, Luke Williams, David Erritzoe, Suresh Muthukumaraswamy, Michael Ashton, Adam Bendrioua, Okdeep Kaur, Samuel Turton, Matthew M Nour, Camilla M Day, Robert Leech, David Nutt, Robin Carhart-Harris

**Author notes:** **Corresponding author:** Christopher Timmermann. 160 Du Cane Rd, Hammermsith Hospital, Imperial College London, W12 0NN, London.

## Abstract

Studying transitions in and out of the altered state of consciousness caused by intravenous (IV) N,N-Dimethyltryptamine (DMT – a fast-acting tryptamine psychedelic) offers a safe and powerful means of advancing knowledge on the neurobiology of conscious states. Here we sought to investigate the effects of IV DMT on the power spectrum and signal diversity of human brain activity (6 female, 7 male) recorded via multivariate EEG, and plot relationships between subjective experience, brain activity and drug plasma concentrations across time. Compared with placebo, DMT markedly reduced oscillatory power in the *alpha* and *beta* bands and robustly increased spontaneous signal diversity. Time-referenced analyses revealed close relationships between changes in various aspects of subjective experience and changes in brain activity. Importantly, the emergence of oscillatory activity within the delta and theta frequency bands was found to correlate with the peak of the experience, and particularly its eyes-closed visual component. These findings highlight marked changes in oscillatory activity and signal diversity with DMT that parallel broad and specific components of the relevant subjective experience and thus further our understanding of the neurobiological underpinnings of immersive states of consciousness.

## Introduction

N, N, Dimethyltryptamine (DMT) is a naturally-occurring serotonergic psychedelic^1^ that is capable of producing experiences that, in intensity, surpass those associated with standard doses of most orally administered psychedelics and indeed most other categories of psychoactive drugs^2, 3^. The subjective effects of intravenous DMT have a rapid onset that is characterized by unusually vivid visual imagery and somatic effects, which arise within seconds of the injection. At high doses, the experience rapidly progresses into a deep and profound immersion – sometimes described as a ‘breakthrough’. This experience is often characterized by a sense of entering into an entirely ‘*other*’ but no less ‘*real*’ world or dimension^3, 4^. It is not uncommon for people to describe encounters with sentient ‘entities’ or ‘presences’ within this perceived other world^4, 5^ and for the experience to subsequently challenge beliefs about the nature of reality and consciousness.

The phenomenology of the DMT experience suggests it may be an especially powerful scientific tool for illuminating the neurobiology of consciousness. DMT experiences can be said to resemble ‘world-analogue’ experiences (i.e. interior analogues of external worlds) – similar to the dream state^6^. It is logical to presume that conscious processing becomes ‘functionally deafferented’ (i.e. cut-off) from the external sensorium in these states, paralleled by what is presumably an entirely internally generated ‘simulation state’, felt as entry into an entirely other world^5^. The rapid, short-acting and dramatic subjective effects of intravenous DMT therefore render it well-suited for investigating the neurobiology of consciousness with functional brain imaging – and this is what we sought to exploit here.

Previous EEG and magnetoencephalography (MEG) studies of psychedelic-induced changes in brain activity have yielded generally consistent results. Broadband decreases in (absolute) oscillatory power have been seen with psilocybin^7^, LSD^8^, and (the DMT-containing ceremonial brew) ayahuasca^9^. Brain imaging measures of the complexity, diversity, or entropy of brain activity have been used to index the quality of a range of different conscious states^10–12^ and MEG studies have shown this to be increased after the administration of a range of psychedelics^13^. Recent advances in analytical methods have allowed for the decomposition of spectral power into its oscillatory and fractal (1/f) components^14^ and these components are thought to have distinct functional relevancies^15^. Moreover, as has been shown to be the case with other atypical states of consciousness^14^ – psychedelics appear to affect 1/f-related activity^16, 17^. Here, we sought to disentangle the specific relationships between different spectral components and different aspects of the DMT experience. Considering the fast-acting effects of DMT and the well-known effect that psychedelics have on vascular activity^1^ (which may be confounded with neuronal activity when using functional Magnetic Resonance Imaging; fMRI^18^), we opted to use the EEG as technique which has optimal temporal resolution and is a more direct measure of neural activity.

The primary aim of our study was to determine the effects of a bolus intravenous injection of DMT (versus a bolus intravenous injection of saline) on the power spectrum and signal diversity of EEG recorded brain activity. Further, we aimed to establish the relationship between these brain activity measures, the real-time progression of the subjective experience and parallel changes in plasma levels of DMT. Finally, both conventional psychometric analyses and more temporally finessed methods – inspired by neurophenomenology^19^ – were utilized to assist the process of mapping between brain and experience. Our primary hypothesis was that DMT would decrease oscillatory power in the *alpha* band and increase cortical signal diversity and that these effects would correlate with changes in conscious experience across time.

## Results

### Subjective effects

Participants were asked to provide ratings of the subjective intensity of the drug effects at every minute for a total of 20 minutes after DMT and placebo administration. Figure 1A displays the group-averaged intensity plots for each minute for a total of 20 minutes post-injection. Paired T-tests revealed that the subjective intensity of the experience remained significantly higher under DMT vs placebo for 17 minutes post-dosing (False Discovery Rate – FDR corrected) and peak subjective effects occurred 2-3 minutes post-injection.

**Figure 1.**
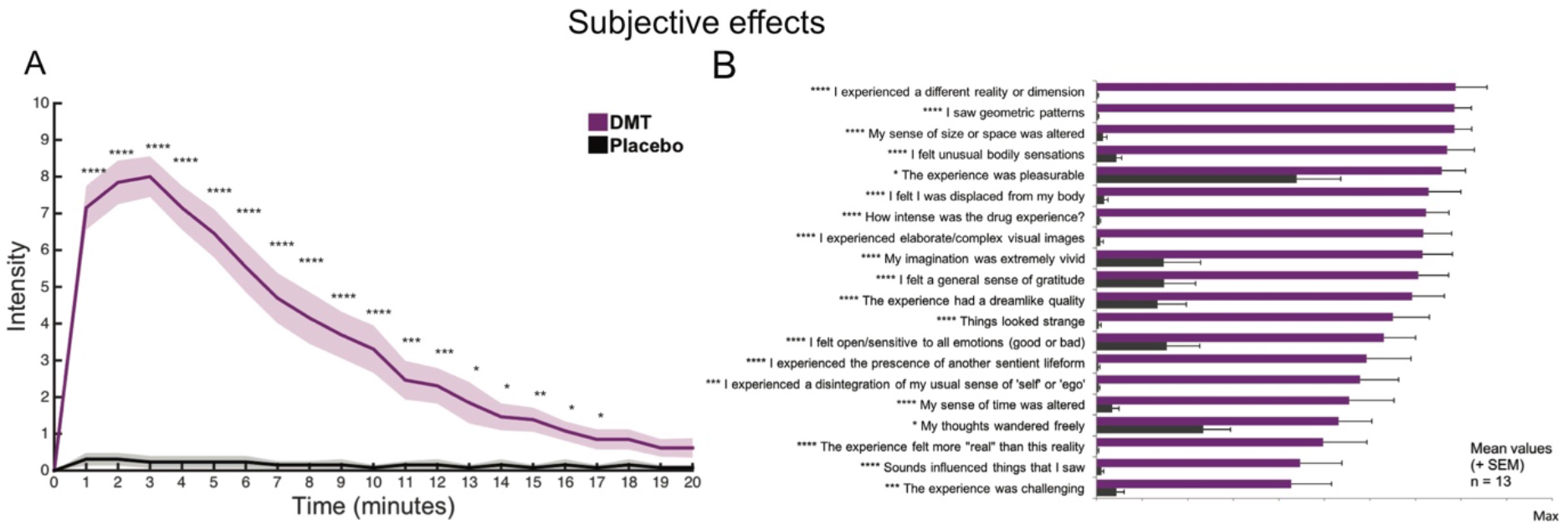
Subjective effects. (A) Real-time subjective effects of DMT and placebo (mean ± SEM) (****p < 0.001; ***p < 0.005; **p < 0.01; * p<0.05, FDR corrected, N=13). (B) Visual analogue scales depict the phenomenological features of DMT and placebo (mean + SEM) (****p < 0.001; ***p < 0.005; **p < 0.01; * p<0.05, FDR corrected, N=13).

Supplementing these basic intensity ratings, participants were asked to rate different aspects of their experiences using various visual analogue scales (VAS). All items were rated significantly higher in the DMT condition compared with placebo (FDR corrected) (Figure 1B). Ratings were given retrospectively, at ~30 minutes following administration, i.e. once the acute effects of DMT had sufficiently subsided.

### Time-averaged EEG results

EEG analyses were done contrasting the group-averaged first 5 minutes of resting state activity following administration of DMT or placebo, focusing on changes in the power spectrum and spontaneous signal diversity (LZs and LZs_N_). One participant was excluded due to excessive movement artifacts following DMT administration. Supporting one aspect of our primary hypothesis, DMT vs placebo contrasts revealed spatially-widespread and statistically-marked decreases in the *alpha band* (max. t(11)=−3.87, cluster p=5.33e-04) and more modest decreases in the *beta* band (max. t(11)=−3.30, cluster p=0.033) (Figure 2A).

**Figure 2.**
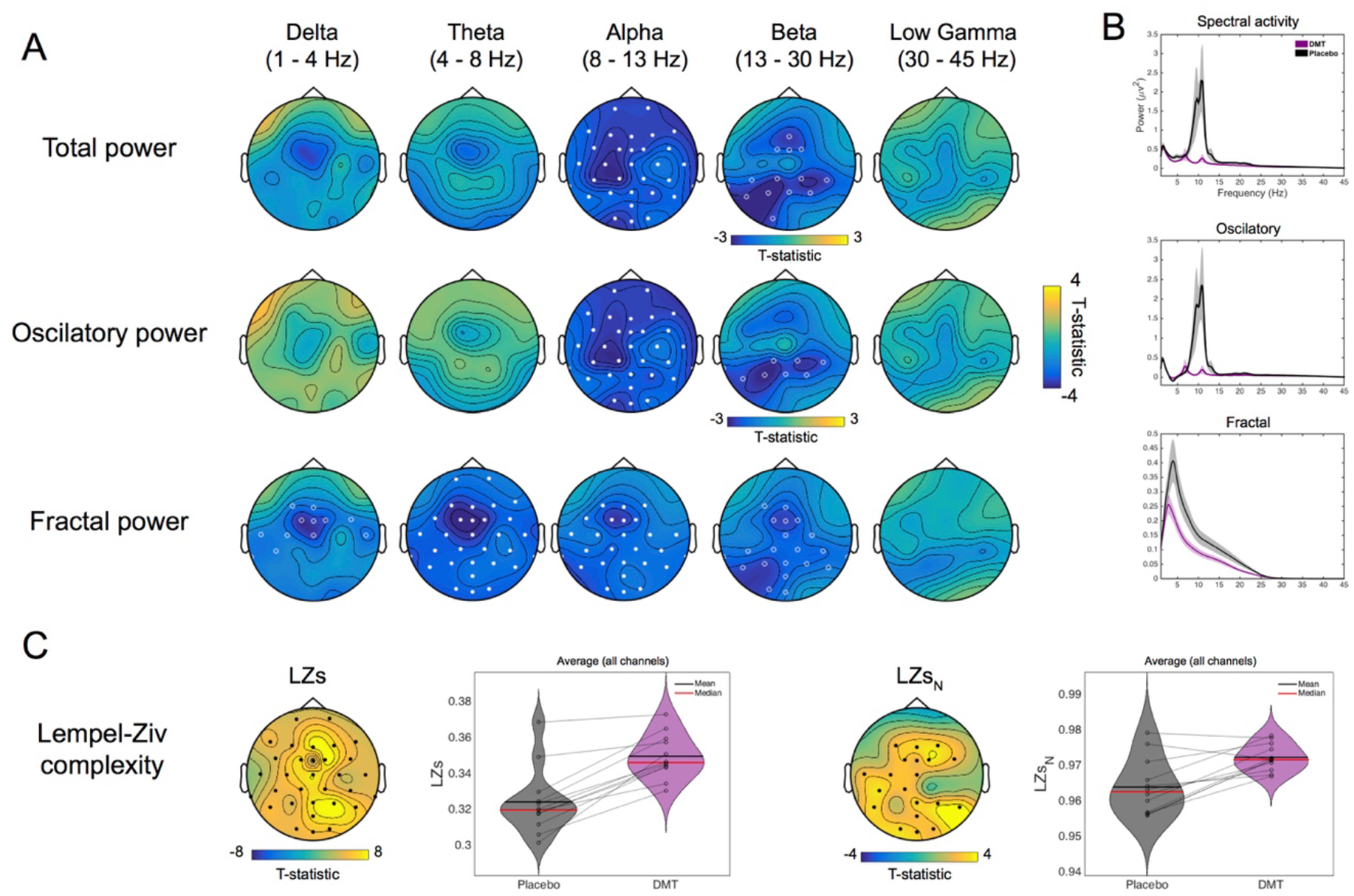
Time-averaged EEG results. (A) The comparison of DMT versus placebo for changes in spectral activity reveals significant decreases for the *alpha* and *beta* bands for conventional spectral power. The decomposed spectra into oscillatory and fractal power, revealed similar results for the first and reductions were seen on all bands <30 Hz for the latter. Increases are seen for both measures of spontaneous signal diversity (LZs and LZs_N_). Filled circles correspond to clusters p<0.01 and hollow circles for clusters p<0.05, N=12. (B) Grand-average spectral power for DMT and placebo corresponding to spectral, oscillatory and fractal (1/f) components of the signal (N=12). (C) Differences (DMT – Placebo) between averaged activity across all channels for the EEG measures displaying the largest effects for each participant before and after injection reveal the decreases in spectral activity and increases in signal diversity for most subjects (N=12) (LZs = Lempel-Ziv complexity, LZs_N_ = normalized LZs).

Decomposition analysis of the EEG spectra into oscillatory and fractal (1/f) components aids in separating the contribution of each of these components to EEG activity, which is useful because unlike fractal activity, oscillatory power has well-known neuronal generators and functional relevance. Results for oscillatory power were consistent with those reported above (*alpha*: max. t(11)= −3.87, cluster p=2.67e-04. *Beta*: max. t(11)= −3.11, cluster p=0.04), while the fractal component showed significantly reduced power within all frequency bands <30 Hz (*delta*: max. t(11)= −3.91, cluster p=0.024. *Theta*: max. t(11)= −4.28, cluster p=0.007. *Alpha*: max. t(11)= −3.67, cluster p=0.014. *Beta*: max. t(11)=−3.48, cluster p=0.02) (Figure 2A).

Signal diversity, as measured by Lempel-Ziv complexity (LZs) and normalized Lempel-Ziv complexity (LZs_N_), was significantly increased under DMT relative to placebo (max. t(11)=8.09, cluster p=2.67e-04; max. t(11)=4.26, cluster p=0.0029), respectively (Figure 2C) – which was consistent with our primary hypothesis.

Results also revealed the emergence of prominent *theta* oscillations under DMT, to the extent that this rhythm became the peak frequency (i.e. the most pronounced rhythm in terms of power) within the 4 – 45 Hz range (mean=7.36 Hz, SEM=0.68) replacing the usually dominant alpha peak (Figure 2B). In contrast, the usual dominant *alpha* peak was maintained throughout the placebo session; mean = 9.28 Hz, SEM=0.62) (t(11) = −2.52, p = 0.029) (Figure 2C). The prominence of this emergent theta rhythm was more evident in the oscillatory power spectrum, i.e. once the fractal component had been removed (Figure 2B and see Fig. S1 for detailed results for DMT and placebo separately).

### Time-sensitive EEG results

In addition to consistent alpha and *beta* reductions, minute-by-minute analysis of the previous results revealed decreases in *delta* and *theta* bands for the first minute only – after which recovery (and increases in *theta* for the oscillatory component) were identified at minutes 2-3 post DMT injection. These results indicate that DMT induces a general decrease in total power across all frequency bands between 1 and 30 Hz. However, there is a transient normalization/increases in *theta and delta* frequencies at the time of peak subjective intensity, which is especially evident in the oscillatory component of the signal. The spontaneous signal diversity measure, LZs, was found to be consistently increased for the whole of the post-injection period, and increases in LZs_N_ were evident from the time of peak intensity onwards (Figure S2).

### Subjective vs EEG effects across time

In order to assess the relationship between the subjective and EEG changes across time, data was segmented into one minute blocks. Results revealed a negative correlation between changes in total *alpha* power under DMT and subjective *intensity* that was significant for all recorded channels (max. t(11)=−16.05, cluster p=2.67e-04)). Decreased *beta* power similarly correlated with higher intensity ratings (max. t(11)=−9.83, cluster p=0.0043). Analysis performed on just the oscillatory component of the signal showed mostly consistent results (*alpha*: max. t(11)= −14.85, cluster p=2.67e-04. *Beta*: max. t(11)= −11.08, cluster p=0.004), although additional positive correlations between *intensity* and *delta* (max. t(11)=4.68, cluster p=0.007) and *theta* power (max. t(11)=7.17, cluster p=0.003) also emerged as statistically significant. Fractal power revealed a negative correlation between (higher) intensity ratings and (reduced) *theta* (max. t(11)=−3.13, cluster p=0.04), *alpha* (max. t(11)=−3.12, cluster p=0.047) and *beta* power (max. t(11)=−5.21, cluster p=0.01). These results reveal the emergence of a functionally-relevant rhythmicity within the delta and theta frequency bands under DMT that was not evident in total power, which includes the fractal component of the signal. In fact, within the theta band, the fractal component appears to behave in an opposite way to the oscillatory component under DMT, i.e. there is increased theta power in the oscillatory component but decreased theta in the fractal component.

Supporting our primary hypothesis, increased signal diversity (LZs) under DMT correlated positively with *intensity* in posterior and central channels (max. t(11)=9.96, cluster p=2.67e-04). After controlling for changes in spectral power (LZs_N_), the relationship between signal diversity and *intensity* remained positive in the posterior channels only (max. t(11)=5.11, cluster p=0.009) (Figure 3).

**Figure 3.**
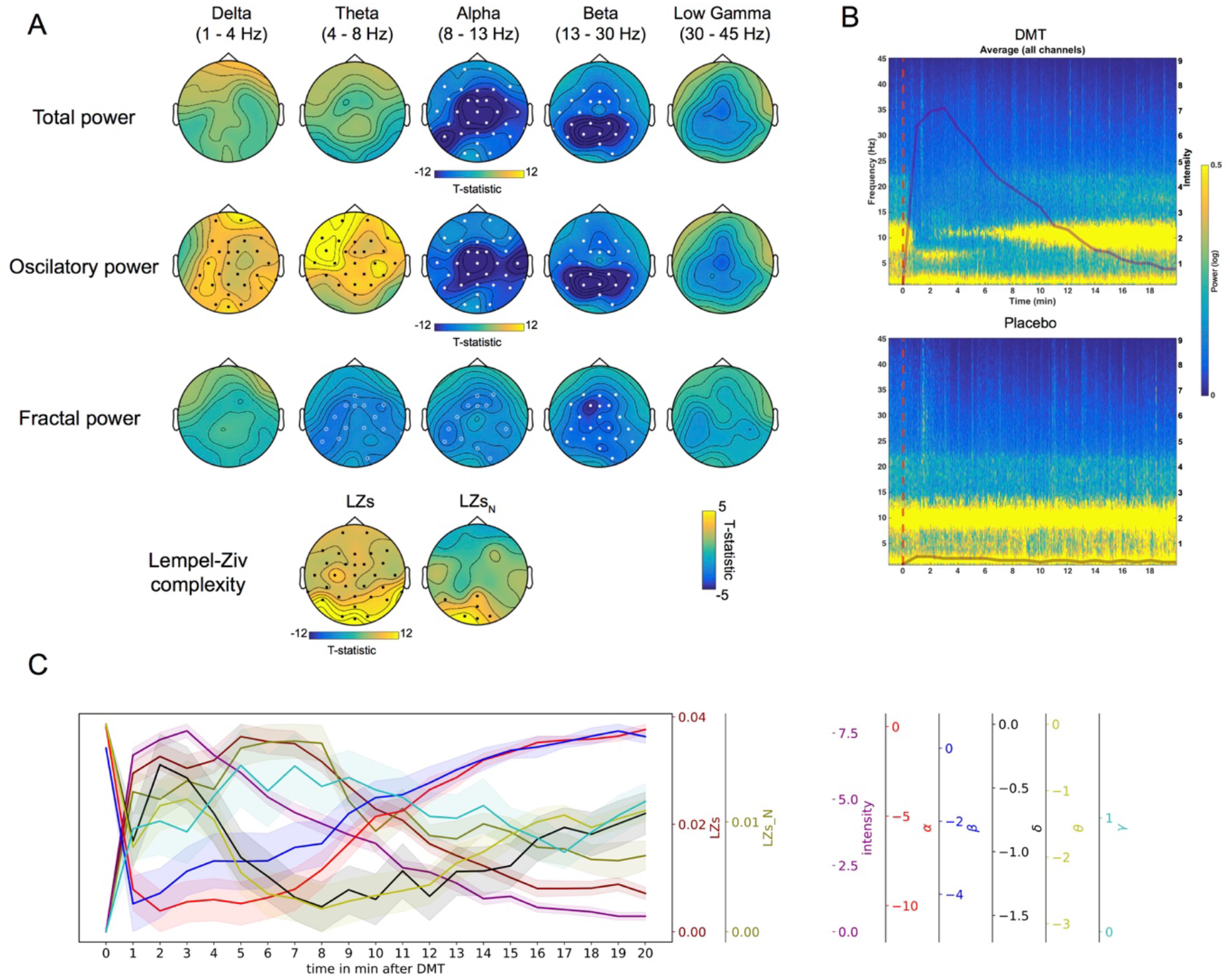
Subjective vs EEG effects across time. (A) Significant inverse relationships were found between real-time intensity ratings and power in *alpha* and *beta* bands for all power measures (including the *theta* band for fractal power). A positive relationship was found between intensity and power at *delta* and *theta* bands for oscillatory power measures of complexity (LZs and LZs_N_). Filled circles correspond to clusters p<0.01 and hollow circles for clusters p<0.05, N=12. (B) Time frequency plot illustrating the associations between intensity ratings and spectral activity for DMT and placebo (red line marks beginning of injection), N=12. (C) Temporal development of intensity, and EEG measures of spectral activity and spontaneous signal diversity (mean ± SEM, N=12). (δ = *delta*, θ = *theta*, γ = *gamma*, α = *alpha*, β = *beta*, LZs = Lempel-Ziv complexity, LZs_N_ = normalized LZs).

### Plasma DMT vs EEG effects

Here we assessed the relationship between changes in the EEG data (frequency bands and signal diversity) and plasma concentrations of DMT across time. In a similar manner to the subjective values, higher concentrations of DMT in the blood were associated with greater reductions in *alpha* (max. t(11)=−23.59, cluster p=2.67e-04) and *beta* power (max. t(11)=−12.61, cluster p=0.04). Similar effects were seen when analyses were performed on the oscillatory component of the signal (*alpha*: max. t(11)= −28.32, cluster p=2.67e-04. *Beta*: max. t(11)= −20.62, cluster p=0.022), while the fractal component showed negative correlation between (higher) intensity ratings and (reduced) *theta* power (max. t(11)=−4.11, cluster p=0.038). As predicted, a relationship was also evident between plasma DMT and increased signal diversity (LZs: max. t(11)=25.61, cluster p=2.67e-04 and LZs normalized: max. t(11)=7.63, cluster p=2.67e-04) (Figure 4).

**Figure 4.**
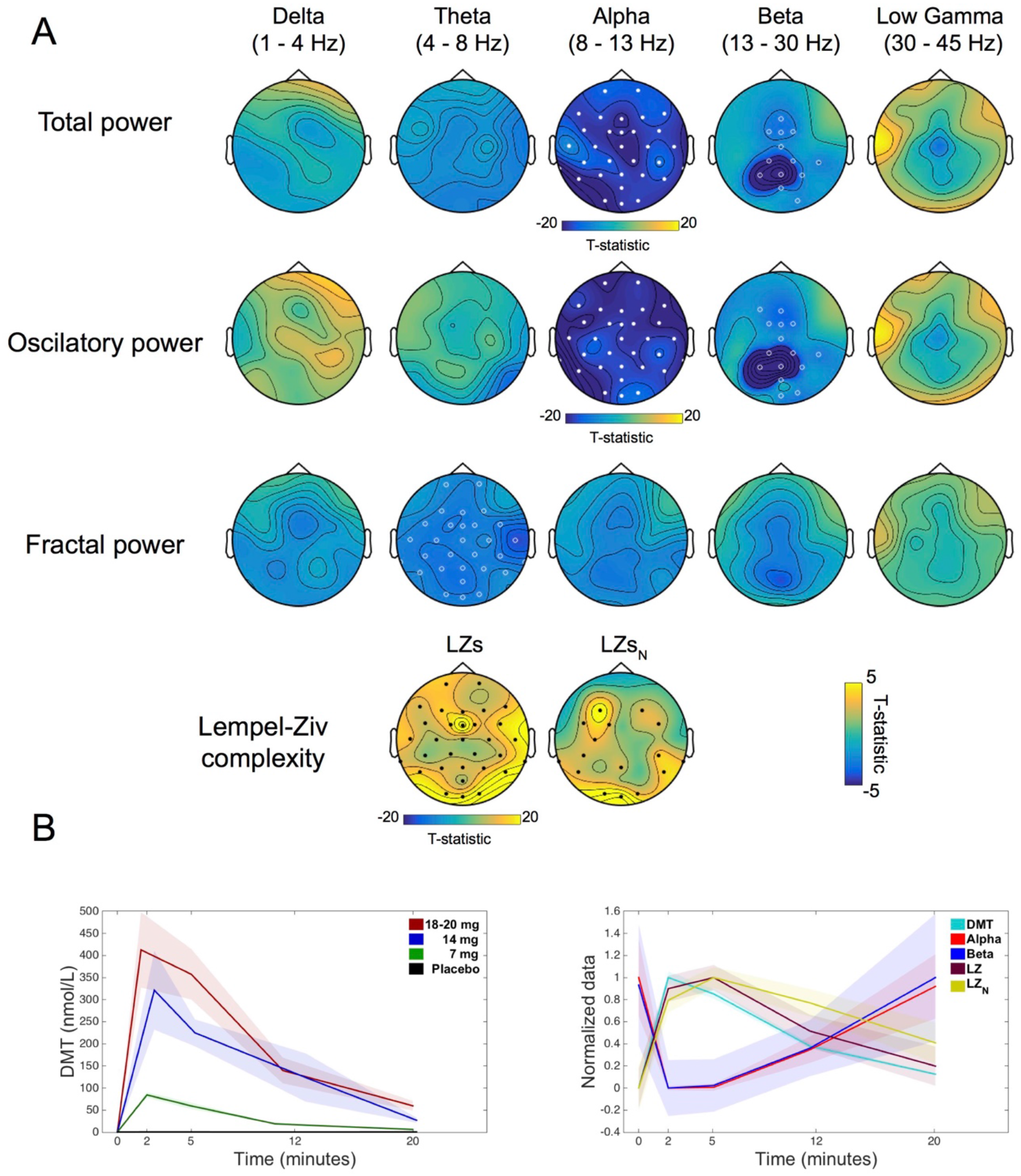
Plasma DMT vs EEG effects. (A) Significant inverse relationships were found between plasma levels of DMT and power in the *alpha* and *beta* bands for spectral and oscillatory power, while the relationship was found for plasma DMT and power in the *theta* band for fractal power. A positive relationship was found between plasma levels of DMT and complexity measures (LZs and LZs_N_). Filled circles correspond to clusters p<0.01 and hollow circles for clusters p<0.05, N=12. (B) Temporal development of DMT plasma concentrations, and EEG measures of total power and spontaneous signal diversity which were found significant to have a significant effect (mean ± SEM, N=12).

### Neurophenomenology

Micro-phenomenological interviews (MPIs)^20–22^ were performed post-hoc to derive distinct components of the subjective experience in a data driven approach – that could then be used to constrain participant ratings referenced to specific time points, i.e. each passing minute. Three major dimensions of experience were found to be common across participants, i.e.: 1) visual, 2) bodily and 3) emotional/metacognitive experiences. These dimensions were extracted from each of the participants’ interviews and then rated by a researcher not involved in EEG analysis with reference to each passing minute within the 20-minute recording period (Figure 5A).

**Figure 5.**
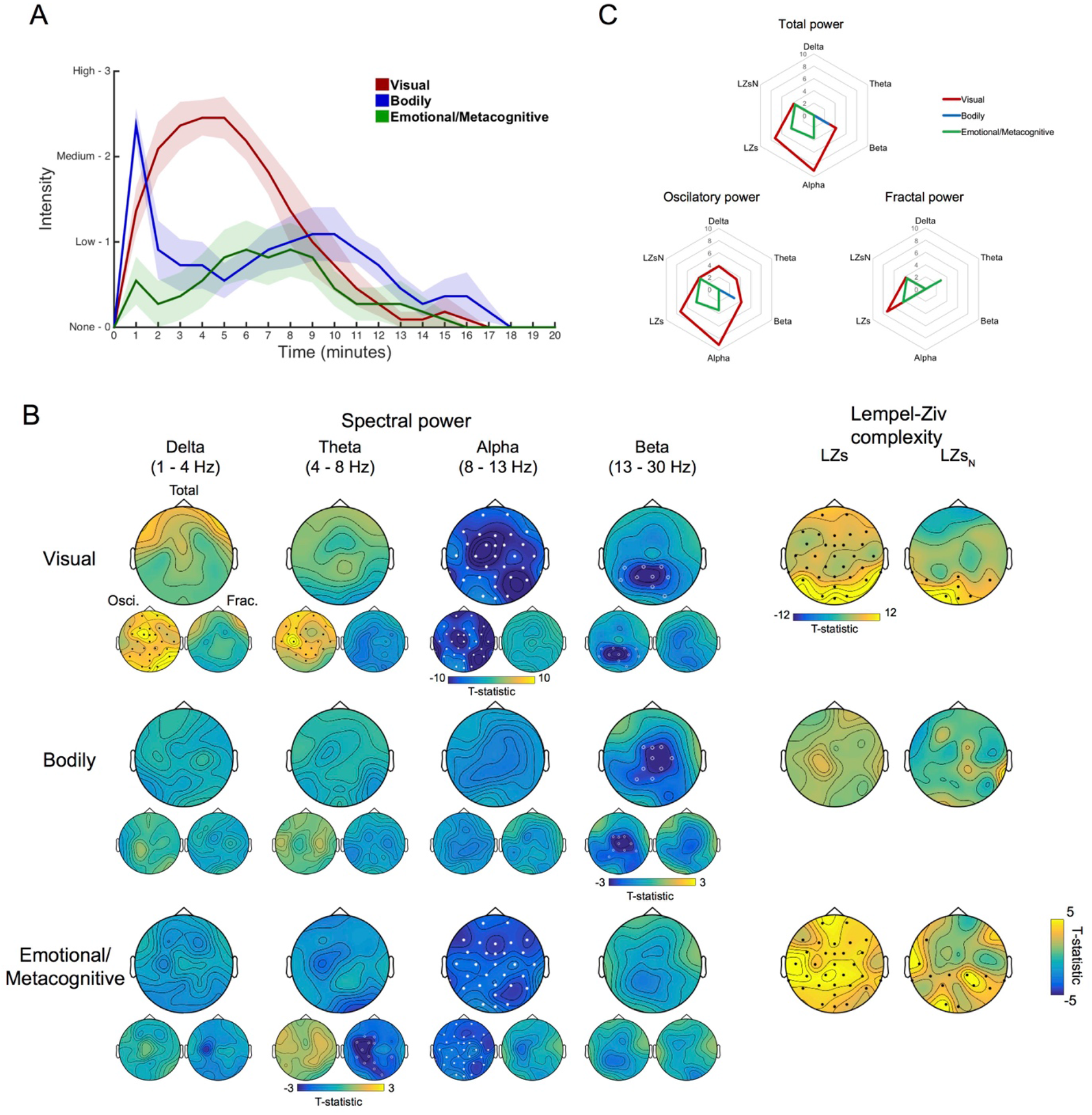
Neurophenomenology. (A) Average ratings (mean ± SEM) regarding the intensity for the three dimensions of experience which were found to be commonly altered across all participants following DMT administration. (B) Significant inverse relationships were found between the progression of visual effects induced by DMT and power at the *alpha* and the *beta* bands, as well as increases in complexity (LZs and LZs_N_). Decreases of central *beta* band power showed a significant association the trajectory of bodily effects. Decreases in *alpha* band power and increases in complexity (LZs and LZs_N_) were significantly associated to the dynamics emotional/metacognitive effects. Mostly consistent results were found with oscillatory power, however an intriguing positive relationship found with power at *delta* and *theta* bands for visual effects and reduced theta activity was linked to emotional/metacognitive effects. Filled circles correspond to clusters p<0.01 and hollow circles for clusters p<0.05, N=11. (C) Radar plots displaying the constellation of EEG effects associated to different dimensions of experience (mean values displayed).

Adopting a data-led approach, we chose to look at components of the EEG that had already shown interesting relationships with intensity ratings, namely: *alpha*, *beta*, *delta* and *theta* power, plus signal diversity measures (LZs and LZs_N_). Results revealed a negative correlation between changes in visual intensity and changes in total *alpha* (max. t(11)= −14.24, cluster p=2.67e-04) and *beta* (max. t(11)= −6.17, cluster p=0.04) power as well as a positive relationship with changes in *delta* (max. t(11)= 6.06, cluster p=0.004) and *theta* (max. t(11)=6.59, cluster p=0.01) power – when just the oscillatory component of the signal was used for analyses. Similar positive relationships between intensity ratings and EEG measures were seen for LZ (i.e. LZs: max. t(11)=16.74, cluster p=2.67e-04 and LZs_N_: max. t(11)=6.62, cluster p=0.008). Ratings of bodily effects were negatively correlated with changes in the *beta* band (max. t(11)=3.17, cluster p=0.0496) only. Lastly changes in the emotional/metacognitive dimension were primarily associated with decreases in the *alpha* band (max. t(11)=−4.56, cluster p=0.008) and increases in the LZ measures (LZs: max. t(11)=6.15, cluster p=0.003 and LZs_N_: max. t(11)=5.39, cluster p=2.67e-04) (Figure 5B). These results support and extend on our primary hypothesis.

### Psychometric correlational analyses

In order to further test the relationship between specific EEG measures and the DMT experience, we performed additional post-hoc ‘subjective rating vs EEG’ correlations using both time-sensitive (minute-by-minute). Time-sensitive analysis revealed relationships that were broadly consistent with those reported above, i.e. negative correlations were evident between (higher) VAS item scores and (reduced) *alpha/beta* power and positive correlations were evident between (higher) VAS scores and (increased) LZs/*theta/delta* – during the period of peak subjective intensity (Figure 6A).

**Figure 6.**
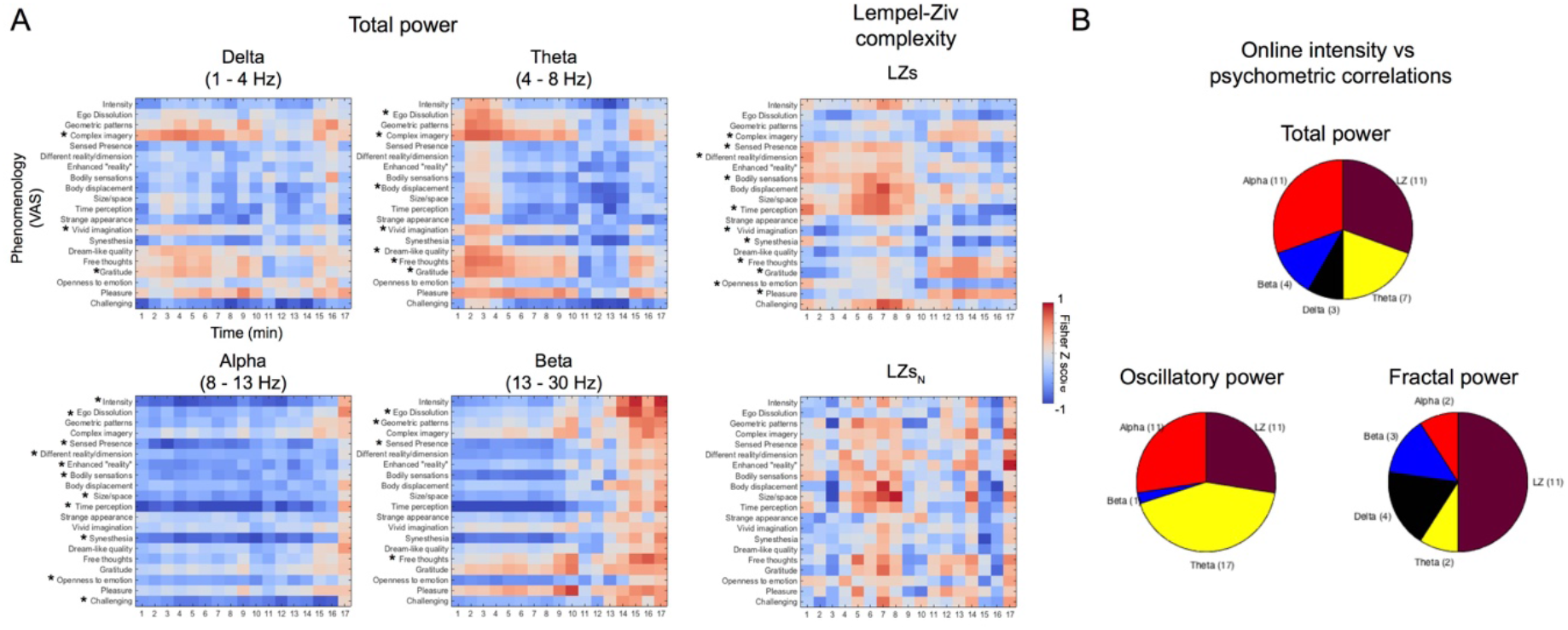
Psychometric correlational analyses. (A) Normalized correlation coefficient values between VAS items and EEG measures for each minute following DMT administration reveal a significant relationship with mean online ratings of intensity. Significant correlations are marked with an asterisk following Bonferroni-correction for multiple comparisons at p<0.05 (See Fig. S3 correlation results with oscillatory and fractal power separately and Fig. S4 for 5-minute averaged data. (B) Pie charts displaying the amount of significant correlations of EEG metrics reveal the prevalence of *alpha* and LZs for total power, the predominance of *theta* for oscillatory power and reduced significance of fractal power across different bands for psychometric correlations.

Lastly, as one might expect, when group-averaged intensity ratings were correlated with Fisher’s Z normalized coefficient scores across time (i.e. minute-by-minute correlational coefficients for VAS item scores vs the relevant EEG-based values) – the subsequent relationships closely replicated those highlighted by the neurophenomenological analyses. Specifically, changes in *alpha* and LZs correlated strongly with the subjective experience when total power was used, while *theta* changes related strongly when just the oscillatory component was used. Finally, after the fractal component was used – only LZs correlated in any notable way with subjective experience (Figure 6B), i.e. the fractal component appeared to be less functionally relevant than the oscillatory component (see Fig. S4 for time-averaged psychometric correlations).

## Discussion

This paper presents results from the first ever placebo-controlled investigation of the effects of DMT on spontaneous human brain activity. Immersion into the DMT state was accompanied by marked decreases in total spectral power in *alpha* and *beta* bands paralleled by marked increases in spontaneous signal diversity and the emergence of *theta* and *delta* oscillations during peak effects. These effects correlated significantly with the characteristic visual effects of DMT. The increases in *delta* and *theta* oscillations were most clearly evident when the oscillatory component was extracted from the fractal component, suggesting that the former is the more functionally relevant component of the signal – at least in relation to these lower frequency bands.

One particularly consistent finding in neuroimaging research with psychedelics is decreased *alpha* power^7,8,23^. Alpha is the most prominent rhythm of the resting-brain, particularly in humans, and particularly in adulthood^24^. *Alpha* power has been linked with high-level psychological functioning ^25, 26^, top-down predictive processing^27, 28^ and related feedback connectivity^29^ – all of which have been found to be disrupted under serotonergic psychedelics^30–32^. Serotonin 2A receptor antagonist (ketanserin) pretreatment studies involving both psilocybin^33^ and ayahuasca^23, 34^ have supported the principle that reduced alpha power under psychedelics depends on their ability to activate 5-HT2A receptors. Here we found strong correlations between *alpha* power decreases, minute-by-minute changes in the subjective intensity, and DMT levels in plasma.

The present study’s findings of profound *alpha* suppression, combined with normalized/increased *delta* and *theta* under DMT may relate to the experience of feeling profoundly immersed in an entirely other world under the drug. The emergence of *theta*/*delta* oscillations, particularly in medial temporal lobe sources, has been classically associated with REM sleep dreaming and related ‘visionary’ states^35, 36^. We propose that the observed emergence of theta/delta rhythmicity combined with the characteristic ‘collapse’ of alpha/beta rhythmicity under DMT may relate to the ‘DMT breakthrough experience’ – a perceptual mechanism by which the brain switches from the processing of exogenously incoming information to a state in which processing is endogenously-driven, as in classic REM sleep dreaming^6^. This is further supported by the positive correlation we observed between participants’ ratings of the visual quality of their experiences and increases in theta and delta power – and decreases in alpha, especially evident once the (putatively) more functionally relevant oscillatory component of the EEG signal was isolated from the fractal component. Although speculative, it is intriguing to consider that the emergent theta/delta rhythmicity under DMT may have a deep (e.g. medial temporal lobe) source and reflect the recruitment of a circuitry that has been classically associated with REM-sleep and medial temporal lobe stimulation – both of which are known to feature complex visionary phenomena^35^.

The increases in signal diversity found here, as elsewhere^13^ may be considered the positive complement of reduced *alpha* power and are consistent with the so-called ‘entropic brain hypothesis’ which proposes that within a limited range of states (i.e. within a critical zone) the richness of content of any given conscious state, can be meaningful indexed by the entropy of its underlying spontaneous brain activity^10, 11^. Based on the present study’s results, revealing a strong and comprehensive relationship between spontaneous signal diversity (a measure intimately related to entropy^37^) and the temporal evolution of different aspects of DMT’s subjective effects, we maintain that entropy-related measures are indeed informative indices of the quality of a given state of consciousness^10, 11^. An increasing number of studies have reported increased signal complexity, diversity or entropy under psychedelics using a variety of imaging metrics and psychedelic drugs^12,38,39^. The increases in signal diversity observed here were associated with the perceived intensity of the experience as well as levels of DMT in plasma.

The neurophenomenological approach and psychometric correlations employed here, using real-time measures and micro-phenomenological interviews, may be seen as a positive step towards the integration of neuronal and first-person reports^19, 40^. The relevant analyses and results allowed us to establish robust and specific relationships between subjective effects (in the visual, somatic and metacognitive/affective domains) and different aspects of the EEG data. Combining multimodal brain imaging (such as using simultaneous EEG-fMRI) with such advancements in subjective data analysis may further aid our understanding of the neural correlates of the psychedelic experience – and indeed other interesting conscious states.

Finally, these results may shed light on some of the neural mechanisms associated to reports showing antidepressant effects of DMT and DMT-containing compounds^41, 42^. Increased *alpha* band power and decreased *delta* band power has been found in depressed populations^43^ and signal diversity has been found to index fluctuations in mood^44^ and depression^45^. It is reasonable to consider that the massive changes in these measures induced by DMT may have implications for modelling and perhaps treating psychopathology.

Some limitations of the present study are worth highlighting. Three different doses of DMT were administered to participants which may have added some variance to our results. Nevertheless, some variance in subjective intensity is useful for correlational analyses. The fixed-order design could be considered a limitation. However, previous psychedelic brain imaging work of ours using fixed-order designs^7^ have yielded consistent results with those seen here and may prevent problematic carry-over effects incurred by balanced-order designs, especially when considering the long-lasting psychological effects psychedelics have been shown to have^46^.

This is the first report on the resting-state brain effects of intravenous DMT in humans. EEG recordings revealed decreased spectral power in the *alpha*/*beta* bands, accompanied by widespread increases in signal diversity. The temporal dynamics of these brain changes closely mirrored the subjective intensity of DMT’s effects. An intriguing finding was the observation of an emergent *delta/theta* rhythmicity during the powerful ‘breakthrough’ state characteristic of high dose DMT, which was mirrored in time by complex visionary experiences. Further work is now needed to more closely scrutinize this feature of apparent order amidst the background of disorder that has more traditionally been known to characterize the psychedelic state^10, 11^.

The present study’s findings significantly advance our understanding of the brain basis of one of the most unusual and intense altered states of consciousness known – a state that has previously been likened to the near-death experience^47^. These findings therefore tell us something important about the neuronal underpinnings of normal consciousness itself – as we can observe what is lost and gained when it transitions in an extreme way – but without the loss of content or awareness. These results may also inform on the nature of analogous states such as dreaming^35, 48^ and the experience of dying^47^ – and in so doing, advance our appreciation of mind-brain relationships in the broadest range of contexts.

## Methods

### Participants and Experimental Procedure

Thirteen healthy participants (6 female, 7 male, mean age, 34.4, SD, 9.1 years) were recruited and provided written informed consent for participation in the study, which was approved by the National Research Ethics (NRES) Committee London – Brent and the Health Research Authority. This study was conducted under the guidelines of the revised Declaration of Helsinski (2000), the International Committee on Harmonisation Good Clinical Practices guidelines, and the National Health Service Research Governance Framework. Imperial College London sponsored the research, which was conducted under a Home Office license for research with Schedule 1 drugs.

Physical and mental health screening consisted in routine physical examination, electrocardiogram, blood pressure and pulse, routine blood tests, as well as a psychiatric interview conducted by a medic. Main exclusion criteria consisted in < 18 years of age, having no previous experience with a psychedelic/hallucinogenic drug, personal history of diagnosed psychiatric illness, immediate family history of psychotic disorder, excessive use of alcohol (> 40 units per week) and blood or needle phobia. A urine test for drugs of abuse and pregnancy (where applicable) and a breathalyser test were conducted on each study day prior to drug administration.

Participants were asked to attend 2 experimental sessions at the National Institute of Health Research (NIHR) Imperial Clinical Research Facility (CRF). Participants were asked to rest at a semi-supine position with their eyes closed throughout the duration of the experiment and eye shades were placed to promote this. EEG data was then collected from one minute prior to drug administration and placebo up to 20 minutes after. Each participant received one of four doses of DMT fumarate intravenously (three received 7 mg, four received 14 mg, one received 18 mg and five received 20 mg) in a 2 ml sterile saline solution over 30 seconds, which was then flushed with 5 ml of saline over 15 seconds. Administration of placebo (2 ml sterile saline) followed the same procedure as previous studies using intravenous administration of DMT^49, 50^. In order to ensure familiarity with the research environment and study team were ensured by using a fixed-order, single-blind design (placebo on the first visit and DMT on the second, which took place a week later). A fixed-order design was used as psychedelics have been shown to induce lasting psychological changes^46^.

Following administration, blood samples were taken at selected timepoints (which were consistent across DMT and placebo sessions) via a cannula inserted in participants’ arm in order to determine DMT levels in plasma. Subjective effects were obtained by asking for intensity ratings in real-time, retrospective Visual Analogue Scales and through ‘micro-phenomenological” interviews.

Acquisition of EEG data was done with a 32-channel Brainproducts EEG system (EasycapMR 32) at a sampling rate of 1000Hz. A 0.1 Hz high-pass filter and a 450 Hz anti-aliasing filter was applied. Additional channels were placed for ECG, EMG and EOG activity, with electrodes placed in the chest, the frontalis and temporalis muscles, as well as placed above and below participants’ left eye.

Blood samples of 10 ml were collected in EDTA tubes prior to and at approximately 2, 5, 12, 20, 33 and 60 minutes after DMT administration. Exact sampling times were recorded. After centrifugation at 2500 g for 10 minutes at 4°C, plasma was harvested and frozen at −80°C. Samples were shipped to Gothenburg in dry ice for quantitation of DMT by high-pressure liquid chromatography with tandem masspectrometric detection. In brief, acetonitrile was added to plasma aliquots to precipitate proteins. After centrifugation at 13 000 g for 10 minutes, 5 μl of supernatant was injected onto the system.

In order to determine the specific progression of effects of DMT over time an independent researcher conducted interviews one day after the DMT session based on the method described by Petitmengin^20, 22^. The Microphenomenological Interview is a technique that is thought to be able to reduce subjective biases particularly affecting first-person reports^20, 21^. It is tailored to facilitate the relationship between subjective experience and its neurophysiological counterparts^22^. Three large dimensions of experience that were common to all participants in the study (visual, bodily and emotional/metacognitive effects) were identified by an independent researcher and subsequently, the intensity of each dimension was rated by the interviewer for every minute using a 4-pont likert scale. Ratings were then used for neurophenomenological correlations.

### Analysis

EEG data was preprocessed using Fieldtrip toolbox^51^. Data was band-pass filtered at 1 – 45 Hz and was visually inspected. Data containing gross artefacts (jaw clenches, movement) were removed from further analysis, as well as segments in which ratings of intensity were asked and collected on every minute following drug administration. Independent Component Analysis (ICA) was performed and the components associated to eye movements and muscle activity were removed from the data. Comparable amount of components was removed across placebo (mean = 5.58, SD = 2.01) and DMT (mean=6.2, SD = 1.66) were removed. Alternative ICA cleaning was performed with less components removed (Placebo mean=4.25 SD =0.92; DMT mean =4.9, SD =0.87), revealing comparable results. The clean data was then re-referenced to the average of all channels and segmented in trials lasting 3 seconds.

Spectral and spontaneous signal diversity analysis were performed for 5 minutes-averaged data before and immediately after DMT/Placebo administration (time-averaged results). Additionally, we performed analysis using 1 minute averages throughout the whole 20 minutes of EEG recordings following DMT/Placebo injection (time-sensitive results). Conventional spectral analysis was performed using slepian multitapers with spectral smoothing of +/- 0.5 Hz using the Fieldtrip toolbox^51^. In order to determine the contribution of oscillatory and ‘fractal (1/f) components to spectral power, the signal was decomposed using the Irregularly Resampled AutoSpectral Analysis (IRASA) algorithm, as described by Wen and Liu^14^ (see Supplementary Methods for details). Resulting spectra from both the IRASA algorithm and conventional analysis were divided in the following frequency bands for statistical analysis: Delta (1-4Hz), theta (4-8 Hz), alpha (8-13 Hz), beta (13-30 Hz) and low gamma (30 – 45 Hz). We then computed the spontaneous signal diversity following our previous study^13^, thus obtaining a score for Lempel-Ziv complexity (LZs) and a normalized score for LZs (LZs_N_) in order to ameliorate the effects of spectral power on these results (see Supplementary Methods and Fig. 7 for a schematic on LZs computation).

**Figure 7.**
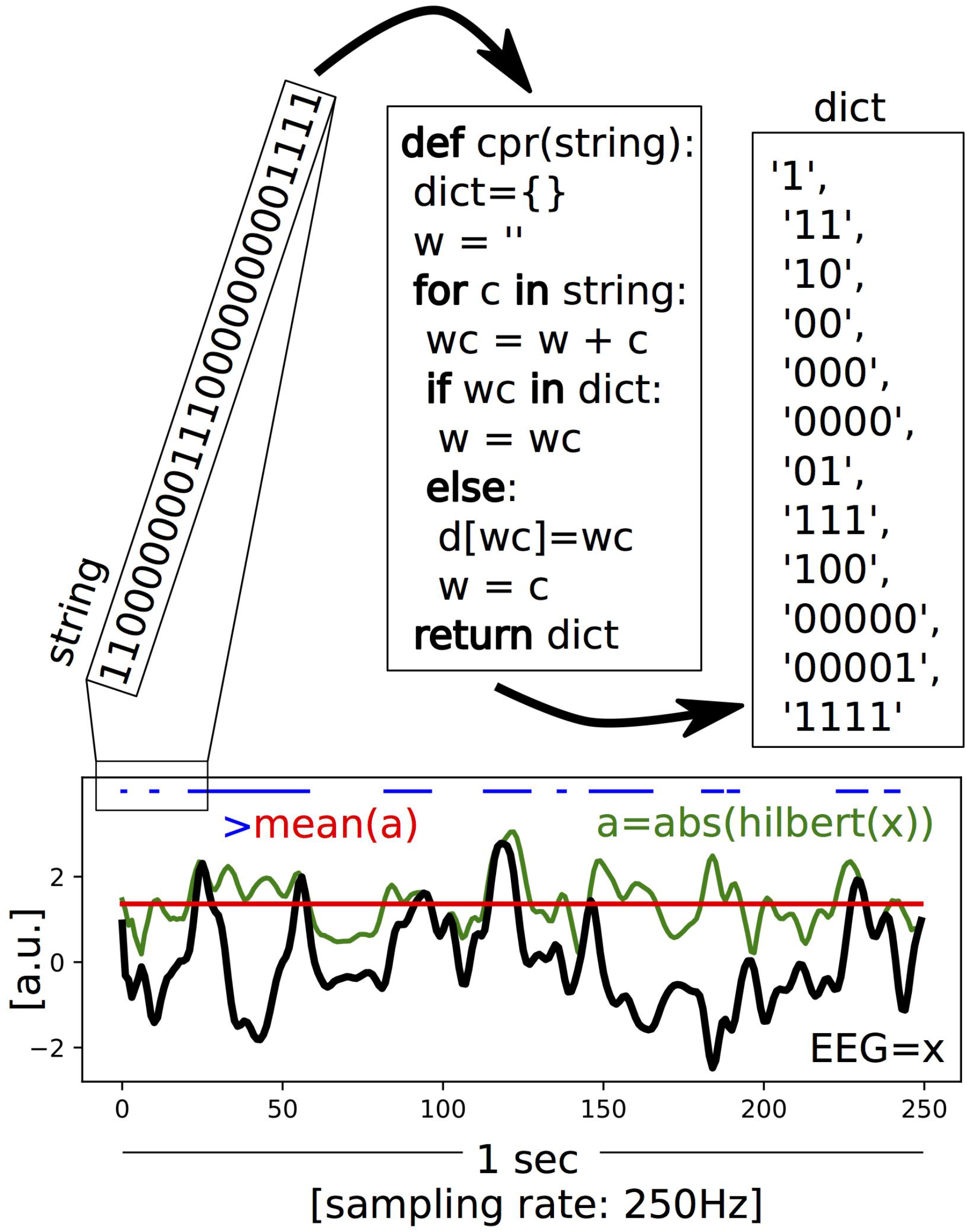
Schematic of the LZs computation. An example EEG signal with a sampling rate of 250 Hz and a length of 1 sec is shown in black (x). The mean (red) of the absolute value of its analytic signal (green, a) is used to binarize the signal (blue). The encoding step of the Lempel-Ziv algorithm is then applied to the first 25 entries of that binarized signal (in this illustration), creating a dictionary of the unique subsequences, which is then normalized by dividing the raw value by those obtained for the same randomly shuffled binary sequence. This provides a value between 0-1 that quantifies the temporal diversity of the EEG signal (LZs).

### Statistical Analysis

Paired T-tests were performed at each time point in order to determine the significance of subjective effects determined by real-time intensity ratings and VAS. EEG-related analysis underwent permutation testing of t-statistics to address differences between EEG and placebo. Correlational analyses were performed between 1) Real-time intensity ratings versus EEG measures across time, 2) Plasma DMT versus EEG measures across time, 3) Scores extracted from microphenomenological interviews and EEG measures across time and 4) VAS and EEG measures (Psychometric correlation analysis). Multiple comparison correction was performed using FDR for all subjective effects measures, and cluster randomization analysis was used to control for multiple comparisons of EEG results with an initial cluster-forming threshold of p = 0.05 repeated for 7500 permutations. (see Supplementary Methods for details).

## Author Contributions

CT designed and led the study, recruited the volunteers, performed the research, analyzed the data, produced all figures (with exception of Figs. 3C, 7 and S1B), wrote and edited the manuscript. LR contributed to study design, helped perform the research and edited the manuscript. MS contributed to data analysis and produced figures 3C, 7 and S1B and edited the manuscript. RM carried out micro-phenomenological interviews and analysis. LW helped perform the research. DE supervised medical aspects regarding the research. SM contributed to data analysis. MA and AB analyzed the plasma samples to detect DMT levels. OK processed the blood samples. ST, MN and CD administered the DMT and provided medical cover. RL advised on study design and contributed to data analysis. DN advised on study design and edited the manuscript. RCH oversaw and designed the study, and wrote and edited the manuscript.

## Acknowledgements

The authors would like to thank all of the study volunteers, as well as Ines Violante, Romy Lorenz and Irina Simanova for advise during data collection and analysis and James Rucker for providing medical cover on 2 occasions This report presents independent research, part of which was carried out at the National Institute of Health (NIHR) Imperial Clinical Research Facility). This work received support from Amanda Feilding and the Beckley Foundation.

## Data Availability

The datasets generated during and/or analysed during the current study are available from the corresponding author on reasonable request.

