## Supplemental Figures and Methods for "Neural correlates of the DMT experience as assessed via multivariate EEG"

### Supplementary Figures

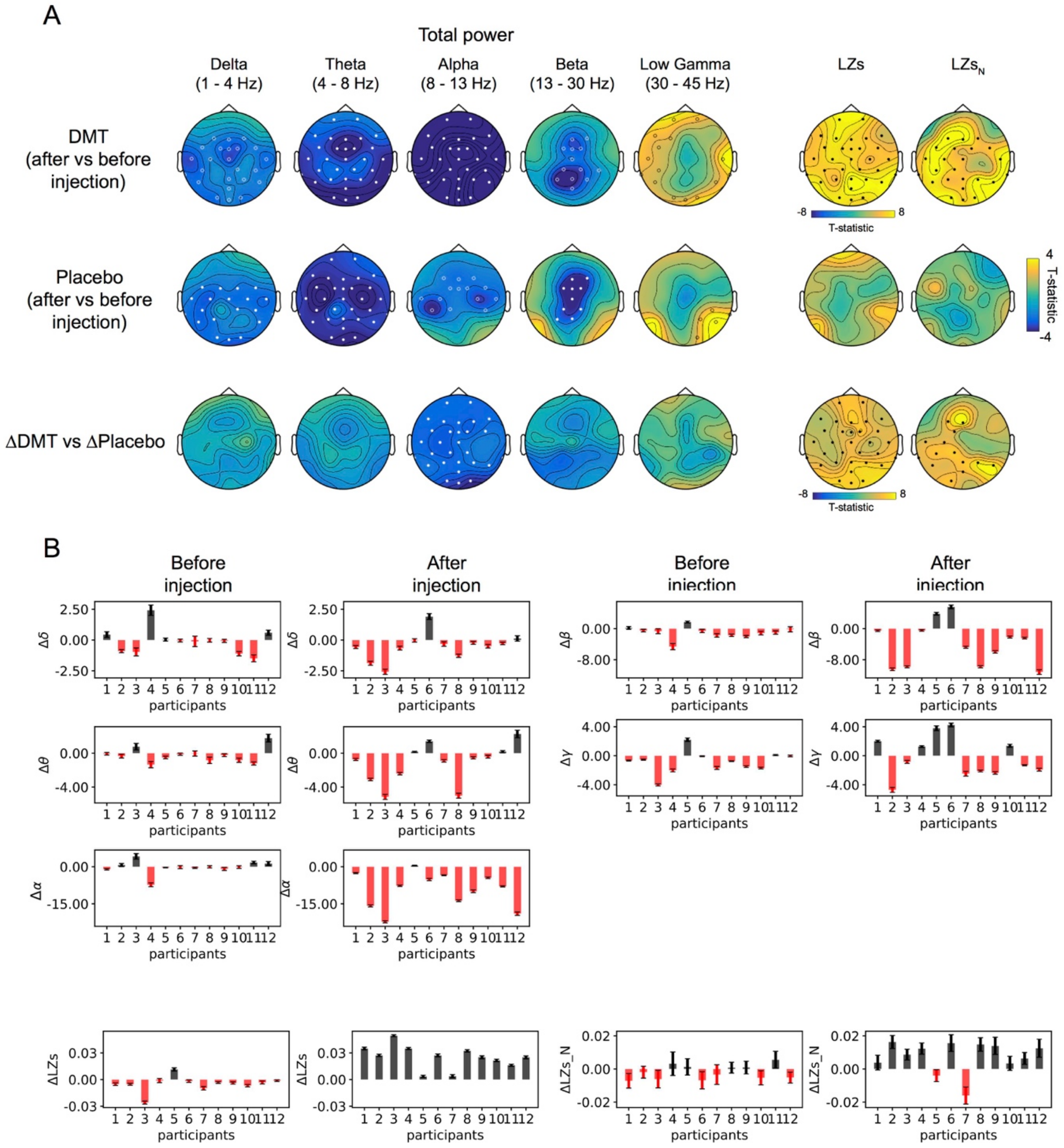


**Supplementary Figure S1. Time-averaged EEG results for DMT and placebo separately.** (A) Significant decreases were found after DMT administration for all frequency bands <30 Hz and increases were found for spontaneous signal diversity. Results from the placebo condition showed reductions in delta, theta, alpha and beta bands, while minor increases were seen for low gamma and LZs. The contrast of DMT versus placebo while taking into account baseline recordings revealed that DMT had an effect on decreases in the alpha band, as well as increases in LZs and LZsN. Filled circles correspond to clusters $p<0.01$ and hollow circles for clusters $p<0.05$, N=12. (B) Differences (DMT-placebo) for averaged activity across all EEG channels at all frequency bands and signal diversity measures before and after injection (δ = delta, θ = theta, γ = gamma, α = alpha, β = beta, LZs = Lempel-Ziv complexity, LZsN = normalized LZs).

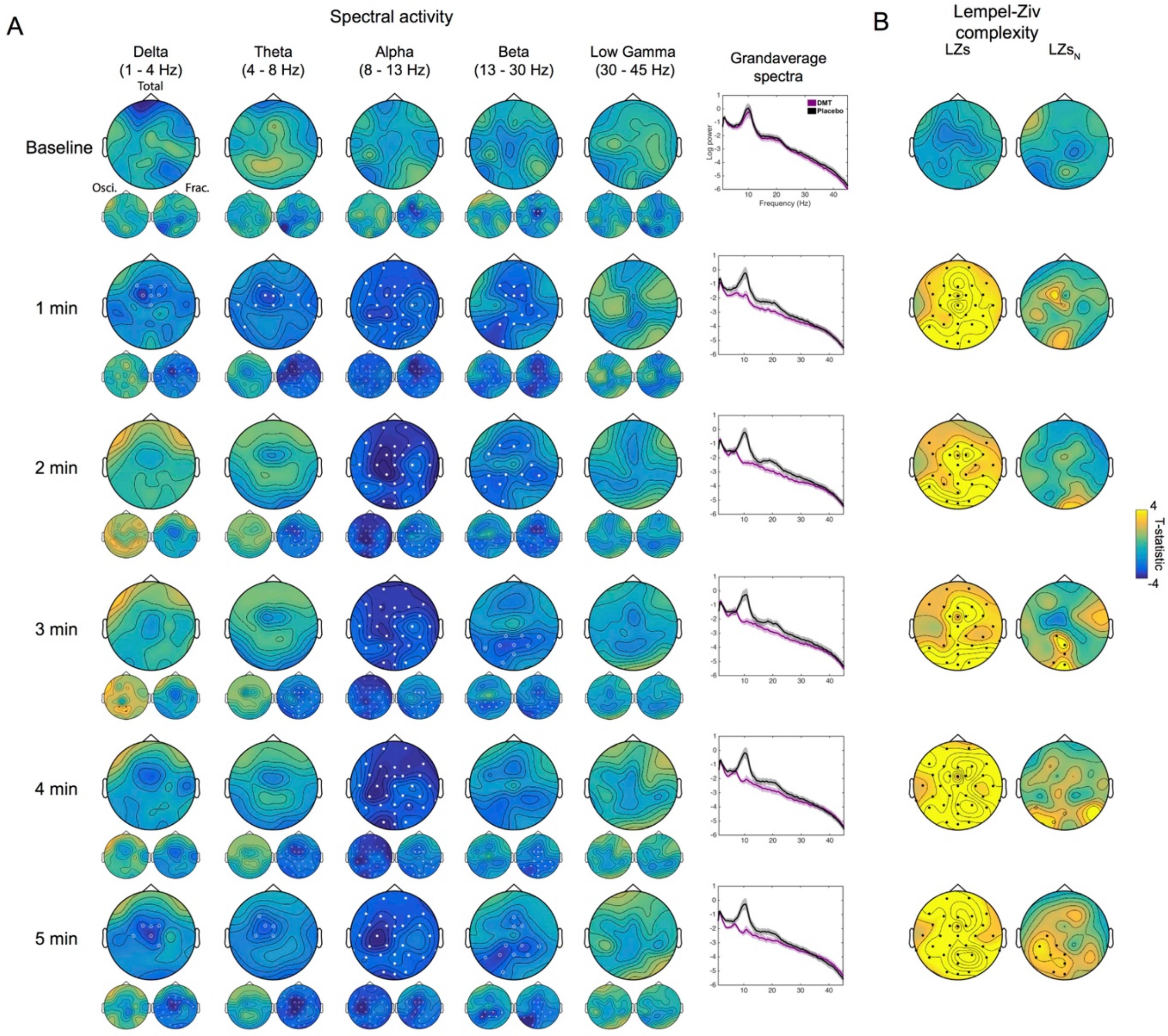


**Supplementary Figure S2. Time-sensitive EEG results.** (A) Results of the comparison of DMT versus placebo using averaged data for each minute, for a total of the baseline and 5 minutes post-injection for spectral activity and the corresponding grandaverage spectral plots (mean ± SEM). Total, oscillatory and fractal power are displayed. (B) Results using complexity measures LZs and $LZs_N$. Filled circles correspond to clusters $p<0.01$ and hollow circles for clusters $p<0.05$, $N=12$. Notice how reductions in lower frequency bands for total power are associated to reductions power in the fractal (and not oscillatory) component.

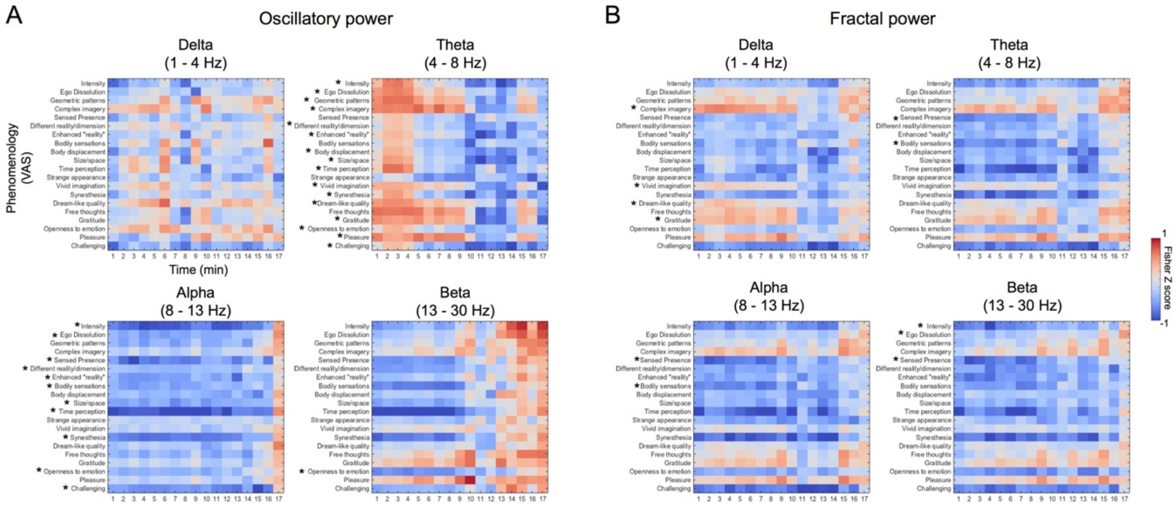


**Supplementary Figure S3. Psychometric correlations for oscillatory and fractal power.** Normalized correlation coefficient values between VAS items and EEG measures for each minute following DMT administration reveal a significant relationship with mean online ratings of intensity for (A) oscillatory and (B) fractal power. Significant correlations are marked with an asterisk following Bonferroni-correction for multiple comparisons at $p<0.05$. Notice the predominance of the *theta* band for psychometric correlations using oscillatory power and the reduced relevance of all bands using fractal power. See Figure 7 for results concerning total power.

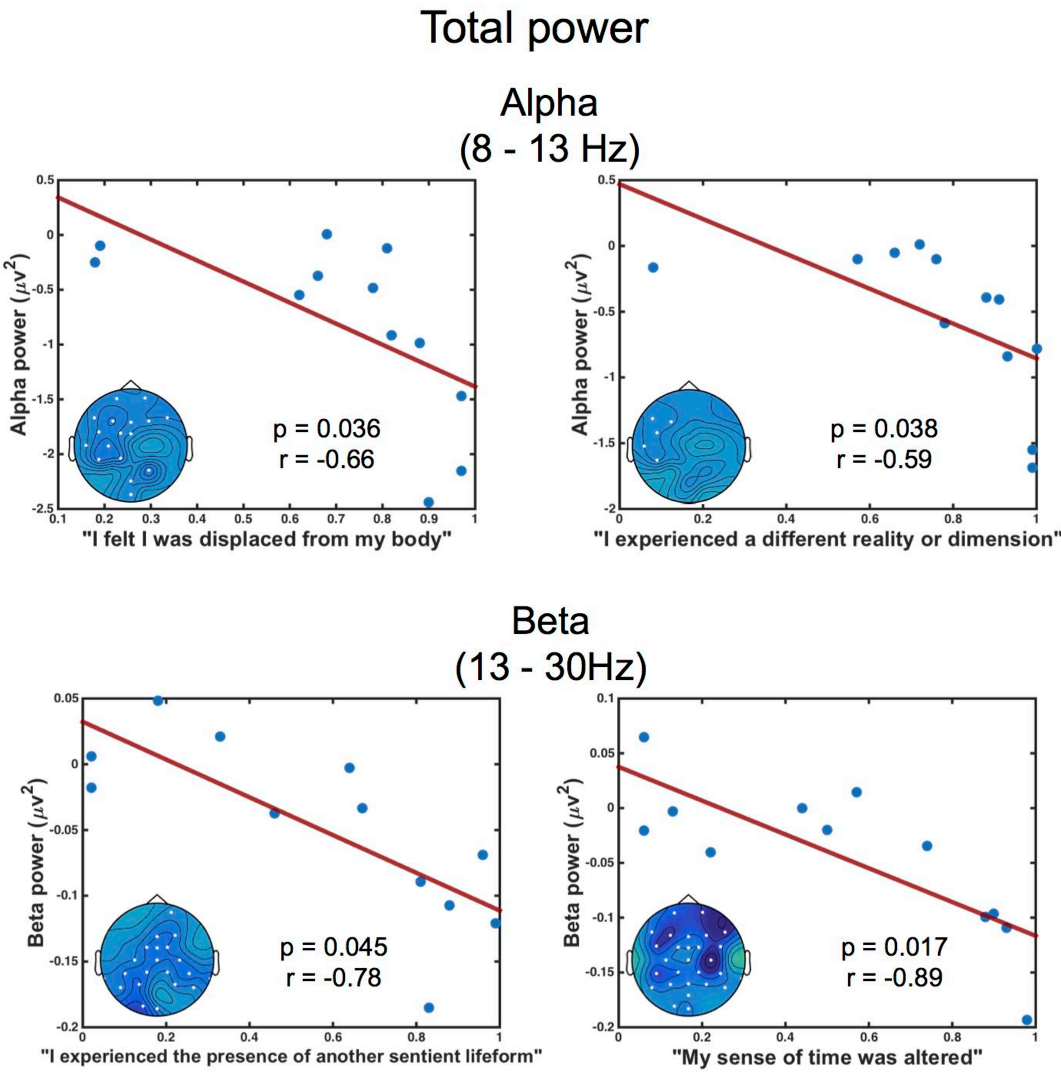


**Supplementary Figure S4. Time-averaged (5-minutes) psychometric correlations.** Significant correlations between power at *alpha*/*beta* bands (total power) and VAS retrospective scores (FDR corrected) (plots display averaged DMT-placebo activity of total power of significant EEG channels).

**Supplementary Methods**

Safety: Ensuring psychological safety is crucial in studies involving psychedelic compounds[1], therefore special attention was given to the physical context and care was placed on the quality of interpersonal interactions between researchers and participants, as these factors are known to influence psychological safety[1,2] and DMT is known for inducing intense emotional effects[3,4]. For this reason, the clinical environment was carefully decorated, maintaining consistency across participants and DMT/placebo sessions. Mood lights, candles and sober decoration was used. When possible medical equipment was covered with blank sheets in order to prevent associations with a sterile, medical context. Additionally, low-volume music was used during the sessions to further provide an experience of comfort and safety and a simple relaxation technique (body scan) was used prior to the administration of DMT and placebo.

Subjective effects: A measure of subjective effects, related to the intensity of drug effects was obtained by asking to rate the intensity of subjective effects from 0 to 10 on each passing minute following drug administration for a total of 20 minutes after injection. Anchors related to the extreme of the scale were explained to participants with a rating of 0 corresponding to “no drug effects” and 10 to “the strongest drug effects imaginable”. Immediately after the effects of DMT subsided and participants were able to do so, (or at ~30 minutes following placebo administration) Visual Analogue Scales (VAS) were completed by participants. Participants were asked to rate each item in relation to the moment in which peak effects of the drug were subjectively felt. The items used were based on effects seen in previous research conducted[5–7] as well as some of the reported characteristic effects DMT[8–10].

IRASA: In order to determine the contribution of oscillatory and ‘fractal (1/f) components to spectral power, the signal was decomposed using the method described by Wen and Liu[11]. The Irregularly Resampled AutoSpectral Analysis (IRASA) algorithm is able to obtain an approximate estimate of the power spectra of the oscillatory component of the signal as follows: Each epoch is segmented in overlapping subepochs which comprise 90% of the signal. The Fast-Fourier Transform (FFT), tapered with a Hanning window is applied obtaining the power spectra for each of the subepochs. Each of the subepochs are then resampled (upsampled and downsampled) by factors of h and 1/h (we used a range of h going from 1.1 and 2.9 using an increment of 0.05 as in Muthukumaraswamy et al.[12]). The power spectra are obtained using the same method as for the non-resampled segments. The geometric mean across the resampled power spectra for each of their corresponding subepochs is obtained which results in an estimation of the "fractal" component. The estimated power of the fractal and original epochs are averaged across all subepochs and the average spectrum of the fractal component is subtracted from the power spectrum of the original signal, which yields an estimate of the oscillatory component for each epoch.

Spontaneous signal diversity analysis: We computed Lempel-Ziv complexity following our previous study[13] (python code https://github.com/mschart/SignalDiversity). The instantaneous amplitude (obtained via Hilbert transform) of each source channel is binarised using its mean over observations as a threshold. I.e. the continuous signal of each source channel is transformed into a string of 3000 binary digits (for our case of 3sec segments at 1000 Hz sampling rate), resulting in a matrix with binary entries with a row for each channel and a column for each observation. To assess the signal diversity across observations, the encoding step of the Lempel-Ziv 1978 (LZ78) compression algorithm (implemented by adapting open source code) is applied to the binary string of each channel. The LZ78 algorithm divides the string into non-overlapping and unique binary substrings. The more diverse the binary string, the more substrings will be listed, with a minimal number of substrings for a sequence of 0s (1s) only. The total number of these substrings for a given channel is what we call Lempel-Ziv complexity (LZs), capturing temporal signal diversity of single channels. We normalize LZs by dividing the raw value by the value obtained for the same binary input sequence randomly shuffled. Since the value of LZs for a binary sequence of fixed length is maximal if the sequence is entirely random, the normalized values indicate the level of signal diversity on a scale from 0 to 1.

In order to test if changes in the diversity measures in the drug vs placebo contrast can be completely explained away by changes in the overall power spectrum, we employed the following control procedure. As in Schartner et al.[13], we assume that effects of the spectral profile on LZs' behavior are multiplicative and thus we divided the LZs score for data by the LZs score of the data after we phase-randomised it. Phase randomisation was performed by first applying a discrete Fourier transform, then randomising phases of the frequencies while keeping their amplitudes fixed, and then applying the inverse Fourier transform (i.e. only the relative temporal positions [phases] of the Fourier sinusoids, whose superposition equals the signal, are randomly changed). We denote Lempel Ziv complexity measures normalised by average scores for phase-randomised data as $LZs_N$. All computations were implemented in python.

Statistical analysis of EEG results: For analysis on time-averaged spectral activity (on the frequency bands reported before) and spontaneous signal diversity, five minute segments of EEG activity before, and 5 minutes and 30 seconds following the end of drug injection were used for each condition. An additional thirty seconds were used for the post-injection data to account for the segments discarded related to the verbal ratings participants were asked on every minute after injection (interruptions lasting ~6 s). Permutation testing of t-statistics were used to address differences between DMT and placebo for time-averaged results. Cluster randomization analysis was used to control for multiple comparisons with an initial cluster-forming threshold of p = 0.05 repeated for 7500 permutations. Additionally, peak frequency in the 4-45 Hz range was calculated for DMT and placebo for the average of all channels and a paired t test was performed. Additionally, DMT vs placebo analyses were performed for averaged EEG activity at each minute following administration (see *Time-sensitive EEG results* in the main article text). Two-tailed hypothesis testing were made for all analyses. The channel displaying the absolute maximum t value is reported and the cluster p value is reported.

Statistical analysis of subjective vs EEG effects across time: For the analyses exploring the relationship between EEG measures and real-time intensity a two-step approach was performed. At the first-level an r value was obtained between the intensity ratings (for the significant 1-17 minutes period after injection) and EEG data corresponding to minute-by-minute averages obtained following

DMT administration. This procedure was repeated for each electrode resulting in an r value per electrode for each participant. Statistical significance was then established at the second level by using permutation testing of t-statistics (as described in the section above) comparing the resulting r values obtained in the previous step against a null condition (in which the r value was replaced by a '0'). The same cluster randomization analysis performed in averaged EEG data was used here to control for multiple comparisons. Two-tailed hypothesis testing were made for all analyses. The channel displaying the absolute maximum t value is reported and the cluster p value is reported.

Statistical analysis of plasma DMT vs EEG analysis*:* The relationship between plasma levels of DMT and EEG activity was obtained using the same two-step method described above, replacing intensity ratings by DMT concentrations in plasma. The segments of EEG data used in this case were one minute averages which were selected in time by keeping a temporal correspondence to each moment in which participants' blood was drawn.

Statistical analysis of neurophenomenology: The relationship between scores extracted from microphenomenological interviews (see Methods in the main text for details) and EEG measures was determined using the same method as real-time intensity, with the difference that ratings of real-time intensity were replaced by ratings for each of the dimensions of experience (visual, bodily and emotional/metacognitive) across time in three separate analyses. Similarly, analyses were limited to the significant 1-17 minute period post-DMT administration.

Statistical analysis of psychometric correlations: In order to address the temporal relationship between subjective effects as determined by VAS and EEG data, post-hoc exploratory correlation analysis was performed using minute–by-minute averages (across all channels) of EEG measures showing significance in previous analyses (*delta, theta, alpha, beta,* LZs and $LZs_N$) and all VAS items. In order to normalize individual differences, we used DMT-placebo data at each corresponding minute (previous time-resolved analyses do not require this step as correlation analysis is done at the single-subject level). These analyses were performed for each minute in the significant 1-17 minutes range post-administration. The resulting values (an r coefficient at each minute and for each VAS item) were normalized using the Fisher's Z transform and subsequently each VAS item was correlated with the average online intensity. In order to increase the specificity of these findings (due to the large amount of significant results) we performed stringent bonferroni-correction across all results in order to determine the amount of VAS items relating to each EEG measure.

Time-averaged psychometric correlations: In addition to minute-by-minute correlation analysis we performed more conventional (but less-sensitive) correlational analysis using 5-minutes averaged EEG data. Correlation analysis were performed between differences between drug conditions for each VAS item and changes seen in *alpha* and *beta* band following injection of DMT and placebo, as well as LZs as these were the measures showing the most consistent effects of DMT, explaining separate contributions of the EEG signal. Correlations were performed on the VAS items related to the characteristic effects specific to DMT[4,9] (i.e. the VAS items "I experienced a different reality or dimension" and "I experienced the presence of another sentient lifeform") and ratings of ego-dissolution. FDR correction was applied and two-tailed hypothesis testing was made.
